# Presence and Perceived Body Orientation Affect the Recall of Out-of-Sight Places in an Immersive Sketching Experiment

**DOI:** 10.1101/2022.10.25.513723

**Authors:** Banafsheh Grochulla, Hanspeter A. Mallot

## Abstract

The orientation of sketch maps of remote but familiar city squares produced from memory has been shown to depend on the distance and airline direction from the production site to the remembered square (position dependent recall, Röhrich, Hardiess, & Mallot, 2014). Here, we present a virtual reality version of the original experiment and additionaly study the role of body orientation. Three main points can be made: First, “immersive sketching” is a novel and useful paradigm in which subjects sketch maps live on paper while being immersed in virtual reality. Second, the original effect of position dependent recall was confirmed, indicating that the sense of presence generated in a virtual environment suffices to bias the imagery of distant places. Finally, the orientation of the produced sketch maps depended also on the body orientation of the subjects. At each production site, body orientation was controlled by varying the position of the life feed in the virtual environment such that subjects had to turn towards the prescribed direction. Position dependent recall is strongest if subjects are aligned with the airline direction to the target and virtually goes away if they turn in the opposite direction.

## 1 Introduction

Imagery of distant places is based on the recall of contents from spatial long-term memory into a working memory stage sometimes called a “representational” memory. Even with a fixed set of spatial knowledge available in long-term memory, the imagined representation of a given target place will differ depending on a number of parameters including the subjects’ imagined heading and view point location as well as their actual position and body orientation at the time when imagination takes place. This has been impressively demonstrated in representational neglect (Bisiach & Luzzatti, 1978; Guariglia, Palermo, Piccardi, Iaria, & Incoccia, 2013), a neurological condition in which subjects are able to recall places from the right side of a familiar city square, but not from the left, where “left” and “right” is defined relative to their currently imagined viewing direction. When the imagined view point changes, the recall pattern changes accordingly.

Representational space can be thought of as a local chart of the imagined environment in which the observer takes a fixed position at the center while remembered objects or elements of the scene are represented at their respective egocentric position (see, for example, Bicanski & Burgess, 2020). Alternatively, it can be conceived of as a graph of known views, each taken from a certain view point in the environment, and connected by descriptors of the egocentric movements required to change from one view point to another (Mallot, Ecke, & Baumann, 2020; Röhrich et al., 2014; Schölkopf & Mallot, 1995). While the general predictions made from both models are largely similar, the view-graph approach seems to lend itself more easily to the generation of pictorial imaginations and allows for anisotropies in representational space.

In behavioral studies with normal subjects, the structure of spatial imagery and representational space has been addressed with various versions of the “judgment of relative direction” paradigm (JRD) as well as with the position dependent recall and production of sketch maps or three-dimensional neighborhood models.

### Judgment of Relative Direction

In the judgment of relative direction (JRD) task, subjects are asked to imagine a known environment from a certain point of view, heading towards some remembered object within this environment. From the imagined heading thus defined, they are then asked to report the relative direction to other remembered objects (Shelton & McNamara, 2001). Recall fidelity depends on the imagined heading direction and is best if it aligns with an intrinsic axis of the imagined environment, in the experiment the long axis of a rectangular room. The intrinsic axis may also be defined by objects placed inside the room. Mou and McNamara (2002) placed several objects in an regular grid and found better JRD performance for mental tasks that involved perspectives that were aligned with one of the two grid axes. On a larger scale, the layout of streets in a city or the corridors in an office building may also provide intrinsic axes of reference (Montello, 1991; Werner & Schmidt, 1999). Reference axes for imagery of spaces on different scales such as an office space within an university campus may coexist and need not be identical (Wang & Brockmole, 2003).

Performance in the JRD task also depends on the subject’s body orientation during recall and is superior if their actual body orientation is aligned with the imagined body orientation or heading (Kelly, Avraamides, & Loomis, 2007; May, 2004; Riecke & McNamara, 2017). This “sensorimotor alignment effect” indicates that despite their engagement in imagery and the JRD task, subjects still maintain a sense of their actual bodily position as might be obtained from path integration and spatial updating. This is true even if both the “actual” and the imagined body pose are defined in a virtual environment only (Marchette, Vass, Ryan, & Epstein, 2014). The representational space generated by imagery of a remote environment thus combines references derived from remembered cues of the imagined space itself (the intrinsic axis) and directional tracking or spatial updating between the imagined and the actual observer position (sensorimotor alignment). For overview, see Julian, Keinath, Marchette, and Epstein (2018); Meilinger and Vosgerau (2010).

### Sketch Maps and Building Tasks

Representational memory is organized along preferred reference axes even if no such axes are explicitly induced by the instruction given to the subject. Basten, Meilinger, and Mallot (2012) have shown that the priming of the recall orientation by imagined travel influences the imagery of distant places. In this study subjects located on an university campus have been asked to imagine walking in the downtown between two familiar city squares, thereby crossing the target square in one of two possible directions. When subsequently asked to produce a sketch map of the target square, the direction of the previously imagined travel primed the recall orientation in the sense that subjects were more likely to sketch the target square as is would have appeared during the imagined walk. The authors conclude that performing an imagined walk activates the representational memory due to automated spatial updating or mental travel or both.

In the Basten et al. (2012) study priming is generated by the activation of visual imagery during imagined travel and may result from a visual memory. Alternatively, it may be a result of a spatial updating process in which memories of nearby locations are automatically activated as soon as the observer approaches that target location. Röhrich et al. (2014) therefore asked passers-by in the city center to produce sketch maps of nearby (walking distance) city squares which were out of sight from the interview location. Sketch maps were rated for orientation and showed significant variation with interview location. Sketch maps of nearby locations are predominantly aligned with the airline direction from the recall position to the target places as if subjects could look through the intermittent buildings. Subjects interviewed at more distant locations (about 2 km away) produced more homogeneous maps showing a standard or “canonical” view of the target independent of the interview location. Here, we use the term “canonical” in analogy of the canonical views known from object recognition (Bülthoff, Edelman, & Tarr, 1995). The imagery of the distant target square thus depends on two directions, first, the airline direction, i.e. on how the square would look could we see through the intermittent buildings, and second, an intrinsic axis of the target square defining its canonical view.

In addition to these dependencies, the bodily orientation of the subjects might also play a role. This was demonstrated by Meilinger, Frankenstein, Simon, Bülthoff, and Bresciani (2016) using a building task instead if freely sketched maps. Subjects were given a set of cards naming popular locations around town and asked to rebuild the configuration. The study confirms the position dependence of the recall orientation and additionally demonstrates an effect of body orientation. Note, however, that the Meilinger et al. (2016) study uses landmark buildings in a larger area, not views of a single city square. The building task was transferred to a virtual environment by Le Vinh, Meert, and Mallot (2020), again with the reconstruction of a specific target square as a task. The study confirmed the overall effect of airline direction; body orientation was not addressed.

### Large-Scale Representational Memory

Unlike object configurations in a room that can be perceived at a single glance, larger areas such as the down-town area of a city have to be explored in a step-by-step fashion, a distinction that has been discussed as vista and navigational spaces by Montello (1993). For the resulting representation, Meilinger (2008) suggested a network of local reference frames or charts, each with its own intrinsic axis. In a simple JRD experiment, only one such chart will be activated and imagery will be based on this chart’s intrinsic axis and the body orientation of the observer. In position dependent recall, however, the actual and the imagined environment are usually part of different local charts each with their own intrinsic axis and the relative position of these charts will affect imagery as an additional factor. In the studies discussed above, this relative position is described by the airline direction and the distance to the target. In the present study, we present novel data on the role of body orientation in position dependent recall.

### VR Methodology

Except for the Le Vinh et al. (2020) study, all studies on position dependent recall discussed above are based on real world experiments. With the present paper, we also want to validate the VR technology as a tool allowing better experimental control. After decades of discussion, it is now generally assumed that virtual environments can provide valid experimental results in spatial cognition, both from non-immersive desk-top setups (e.g. Ruddle & Jones, 2001) and with motion tracking and head-mounted goggles (e.g. Avraamides & Kelly, 2008; Kelly et al., 2007; Marchette et al., 2014). However, some problems remain which are likely due to limitations in the fidelity of visual input and in the feedback from the subjects’ bodily motion, expecially in non-immersive set-ups (Lessels & Ruddle, 2005; Richardson, Montello, & Hegarty, 1999).

One additional problem arising in the transfer of the position dependent recall paradigm in a virtual environment is the task to be performed by the subjects. Le Vinh et al. (2020) used an interactive building task in which a joystick was used to grab building blocks from a reservoir and place them on a work space. Here we explore an alternative approach called immersive sketching, in which the subject uses an ordinary drawing board while still immersed in the virtual environment. The sketching process is recorded with a camera and overlaid to the VR simulation as a life image.

## 2 Methods

### Participants

A total of 100 healthy adults, 43 of them male, aged between 18 and 60, participated in our studies. All included subjects successfully passed a pre-experiment. 80 participants were randomly assigned to 4 groups of 20 people each for experiment one and 2 groups of 10 people each for experiment two. Before the study all the participants confirmed that they had no problem with virtual reality and wearing an Oculus RIFT head mounted display and that they had been living constantly in Tübingen for more than two years.

Participants were informed that they could stop and quit the study any time during the experiment without providing any reason. All participants gave written informed consent and received a payment of 8 Euro for their participation.

### Setup

Virtual environments were presented with an Oculus RIFT head mounted display while subjects were seated on a rotating chair as shown in Figure 1. The chair has been prepared for the experiments in the following way:

1. An additional small desk for carrying a laptop has been attached behind the backrest. In the experiments the subjects were able to rotate on the chair without any distraction with the cables of the video equipment.
2. A second desk is attached in the front of the chair similar to a tablet arm desk. For the experiments a piece of paper is attached on the desk that allows the subjects to draw a sketch map. The paper is fixed with a clip on the desk.
3. A camera stand with a camera is attached to the tablet arm desk. The camera is used to record a live video of the paper fixed on the tablet arm desk. The live video is shown in the Oculus RIFT (using a Unity WebCamTexture) which allows the subjects to draw on the paper while being immersed in the virtual environment.

**Figure 1:**
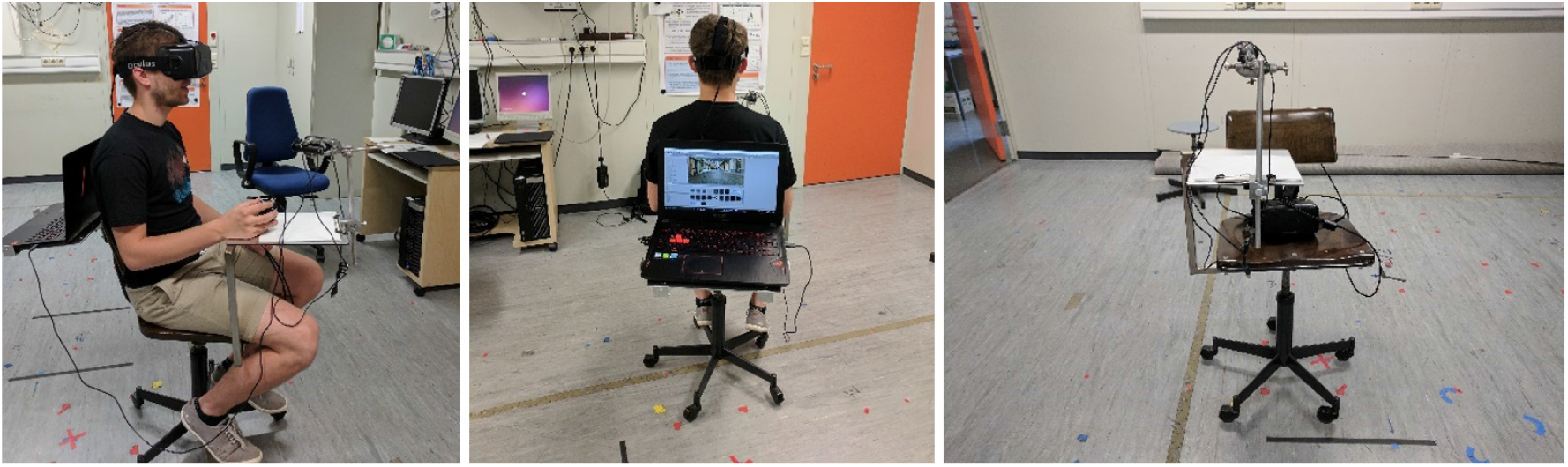
The set up for the experiments consists of a modified rotating chair. Attached to the chair is a small desk for carrying a laptop and a tablet arm desk for drawing the sketch maps. Attached to the tablet arm desk is a camera stand with camera that captures a live video from the sketch paper that is displayed in the virtual environment.

We have used square pieces of paper for the experiments so that the subject is not biased for drawing horizontally or vertically. As the piece of paper was fixed with a clip to the desk it was not possible for the subjects to move or rotate the paper while sketching.

Videos for the pre-experiment were captured on site in Tübingen using a Canon Power Shot G7 camera. They were presented in the Oculus RIFT, which in this case was used just as a simple displaying device, i.e. in open loop.

For the main experiments we have set up a virtual environment in Unity using C#. For each sketching location (S1, S2, S3) we created a 360-degree panorama of the location by taking twelve pictures at 30 degree intervals with the Canon Power Shot G7 camera mounted on a tripod. The twelve pictures were stitched to a cylindrical panorama using Microsoft Image Composite Editor. The three panoramas are shown in Figure 2. All pictures and videos were taken during early morning hours of avoid imaging of passers-by.

**Figure 2:**
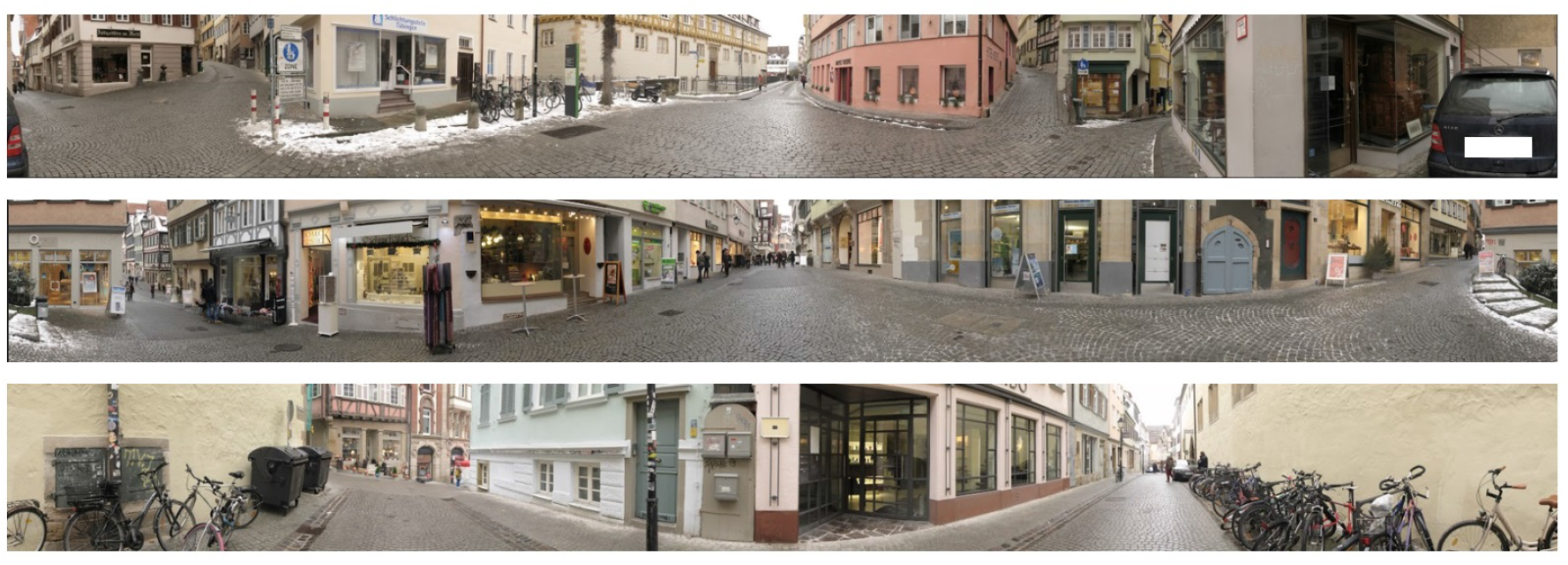
The panoramas of sketching locations S1, S2, and S3.

The sketching locations were modeled by placing the panoramas as textures in the inside of a virtual cylinder. For the top of the cylinder we have used matching images of skies, for example blue sunny sky or cloudy sky. For the bottom of the cylinder we have chosen images of a matching ground, for example cobblestone. The live picture of the sketching paper fixed on the tablet arm desk is displayed in a square work space appearing in the bottom part of the cylinder. This allows the subjects to draw a sketch map on the paper without taking off the goggles. The texture of this square is set to Unity’s WebCamTexture that takes and displays the camera’s live image on the square instead of a static texture or plain color. The work space for sketching is placed either towards or away from the target locations, such that subjects have to produce their sketches while bodily oriented towards or away from the target square.

The subject’s point of view in the experiments was placed in the center of the cylinder. The Oculus RIFT allowed for accurate tracking of the movement of the subject’s head so that the subjects could explore the virtual environment. We have only tracked the rotation of the head; translational movement produced with the upper body while seated was not fed back to the VR simulation. I.e., if the subjects did produce translational movements, the cylinder would virtually move with them and the image remained unchanged.

During the experiment, the experimenter could switch between the three sketching locations using the space bar of a laptop computer, thereby “teleporting” the subjects from one location to another.

### Task

In all experiments, we used three sketching locations (S1, S2, S3) and two target locations (T1: market square, T2: timber market), see Figures 2 and 3. All locations were well-known places in the historic center of Tübingen. All locations were in walking distance of each other but mutually out-of-sight.

**Figure 3:**
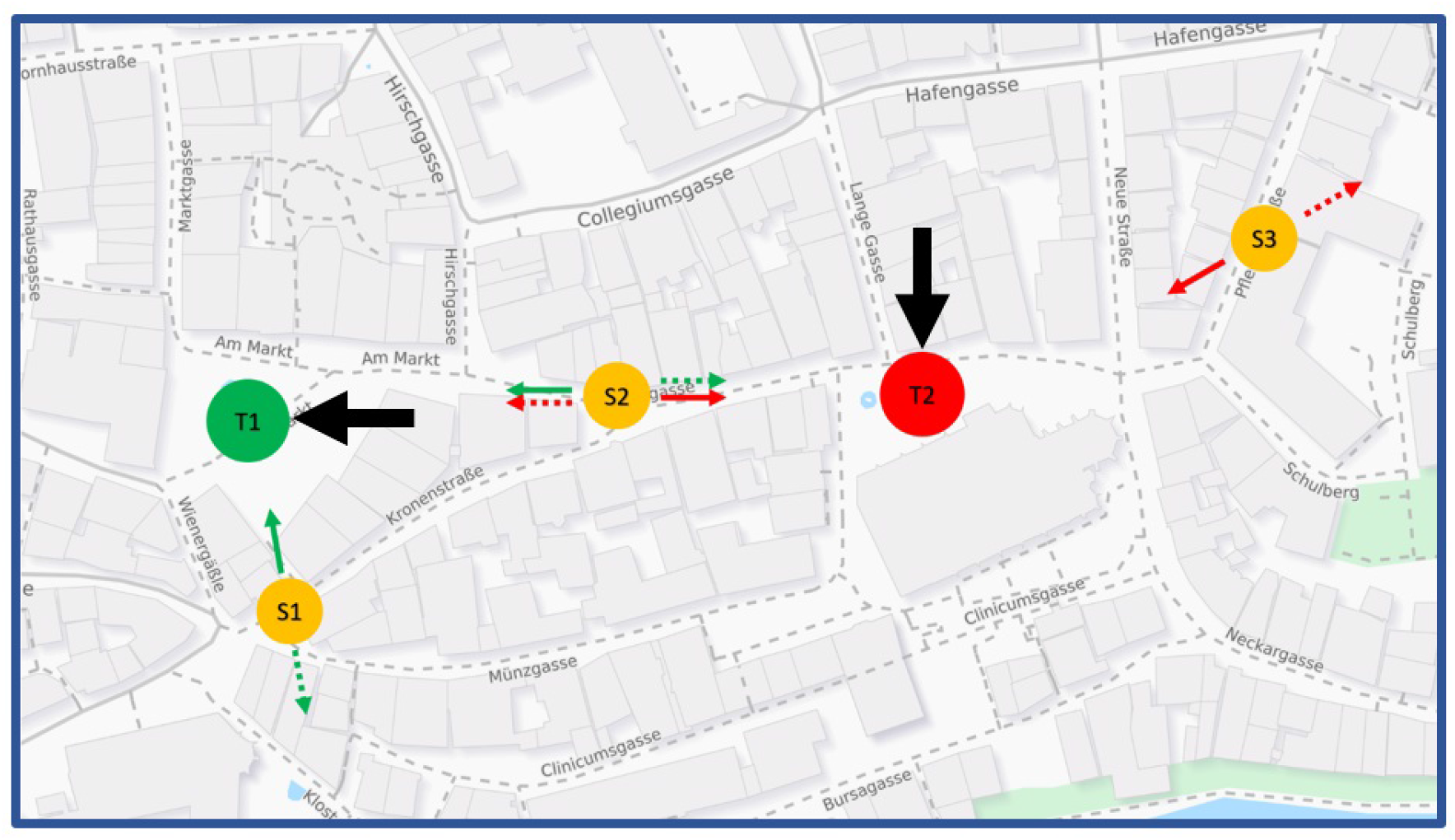
Map of Tübingen city center with the two target locations T1 “market square” and T2 “timber market” and the sketching locations S1, S2, and S3. Red and green arrows: sketching location and body orientation when sketching T1 (green) and T2 (red). Solid arrows mark “towards” condition, dashed arrows mark “away” condition. The heavy black arrows show the canonical viewing direction of the target locations as reported by Röhrich et al. (2014). Map source: www.openstreetmap.org

Each of the two target locations was sketched from two different directions provided by the two closest sketching locations. This results in the four sketching tasks S1-T1 (at S1, sketch T1), S2-T1, S2-T2, and S3-T2, see Figure 3. In addition, we varied the subjects’ body orientation by placing the life image of the sketching paper either towards the street leading from the sketching location directly to the target (“towards condition”) or offset from this direction by 180 degrees (“away condition”). In the “towards” condition, subjects might therefore imagine to just move forward to walk to the target. The “exit direction” is generally similar to the airline direction but slight deviations may occur due to the city street raster. In the “away” condition subjects were required to produce their sketches while imagining a target located behind them.

In total, these variations result in eight different tasks. In order to avoid interactions between repeated sketching tasks, each subject produced two sketches only.

## 3 Procedure

### Pre-experiment

Before the main experiment started, we performed a pre-experiment to test whether the subjects had a good knowledge of the Tübingen city center and were familiar with the locations that we used in our experiments. Subjects were shown three videos exploring the sketching locations S1, S2, S3. For this, the video goggles were used in open loop. Each video started at a small distance from the sketching location and showed a walk through the immediate environment including views into streets connecting to the target locations. However, the target locations themselves were not visible in any of the three videos. After watching the video, the subject answered the questions:”Do you know this place? Do you know what this place is called? When was the last time you visited this area? Why did you visit this area?”

Subjects who failed to name all three sketching locations correctly or reported to have last visited them more than 30 days ago were excluded from the main experiment.

### Experiment 1

After successfully passing the pre-experiment the experimenter explained the experiment to the subject by reading a note about the procedure of the experiment. The subjects received this note also in writing. The experimenter pointed out the equipment including the rotating chair, the camera and the fixed square paper. They clarified that the subjects were allowed to rotate on the chair during the experiment whenever necessary and that they would be asked to draw some layouts on the paper while still wearing the goggles. Subjects were also informed about a relaxation room which they might want to use should they feel sick during the virtual reality experiment.

At the beginning of each sketching task, subjects were teleported to the respective sketching location facing north. They would then explore the environment by looking around (closed loop VR-simulation for rotations only) until they indicated that they had made themselves familiar with the location. The experimenter then asked them to imagine the target location and produce a sketch on the paper provided. To do this, they needed to turn into the direction in which the work area with the live video stream had been provided, either towards or away from the target location.

Each subject completed two sketching tasks, one for each target location. They were randomly assigned to one of four groups of 20 participants, called A-towards, A-away, B-towards, and B-away. The tasks for each group are listed in Table 1. After finishing the first sketching task, the second was started without delay.

**Table 1:**
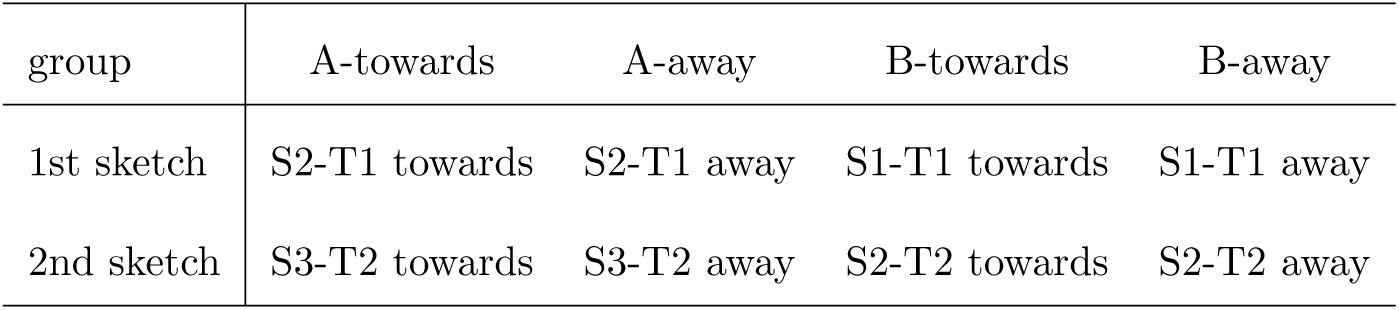
Design of experiment 1. For details see text

### Experiment 2

To understand the effect of immersion and presence on the representational memory, we conducted a second experiment in which sketching was repeated after a delay period and outside the virtual environment. We have tested 20 participants (sex not recorded) who were assigned to two groups C-towards and C-away with 10 subjects per group.

Experiment 2 consisted of two phases: Phase 1 for groups C-towards and C-away is identical to experiment 1 of groups A-towards and A-away, respectively, i.e. subjects perform sketching tasks S2-T1 and S3-T2. After the subjects have finished sketching and the experimenter collected the sketch papers the subjects put away the video goggles and took a 10 minutes break during which they are were asked to stay in the experiment room. They were offered refreshments or might have a look at books available in the room which were irrelevant to the study. However, they were not allowed to browse in the internet or talk on the phone. After the 10 minutes break, phase 2 started in which the subjects were asked to draw the layouts of the same target locations as in phase 1 again, but this time without being immersed in the virtual environment nor with any additional information from the experimenter. Finishing the drawing of the target locations concluded the second phase of the experiment and the experiment itself.

In total each subject produced four sketch maps: two maps of locations T1 and T2 before the break and within the virtual reality environment, and again two maps of the two target location after the break.

## 4 Results

The results from both experiments will be presented together.

Figure 4 shows four examples of the produced sketch maps. The maps have been categorized by two raters according to the cardinal directions north, east, south, and west and intercardinal directions northeast, southeast, southwest, and northwest. For this task the two raters could use any aid necessary, such as their own local knowledge of the area, pictures, or maps. The raters worked independently from each other and agreed on the directions in all cases. Figure 5 shows the distribution of the rating results for sketch maps produced at the three sketching locations S1, S2, and S3, both for the towards and away conditions.

**Figure 4:**
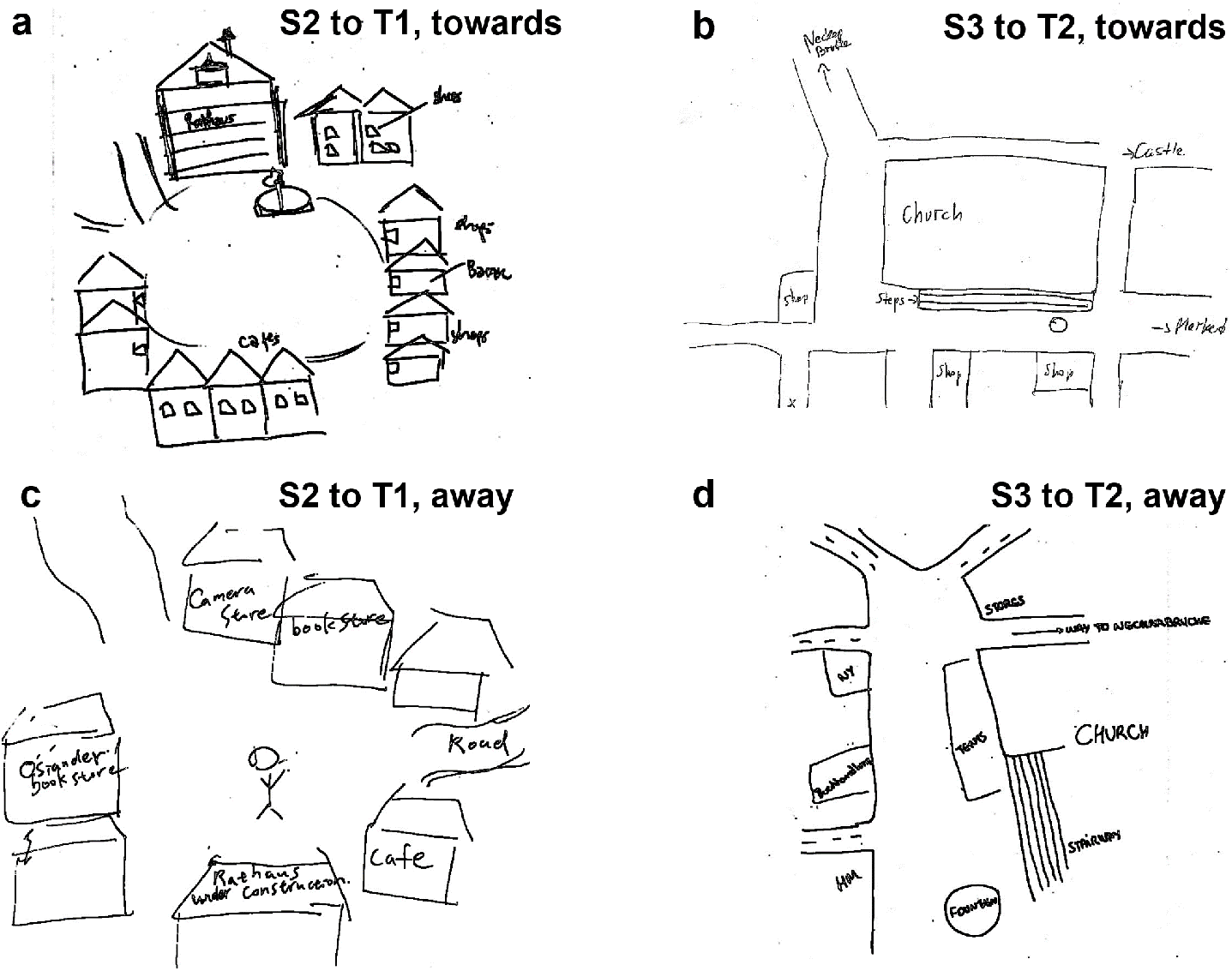
Examples of drawn sketch maps for different target and starting locations and different orientations in the starting locations. **a** and **c** show target location T1, **a** is oriented east (“Rathaus”, City Hall up) and **c** is oriented west (City Hall on bottom). **b**, **d** show target location T2, **b** is oriented north (church up) while **d** is oriented west (church right). Note that the paper orientation while drawing is also defined by the scribbling.

**Figure 5:**
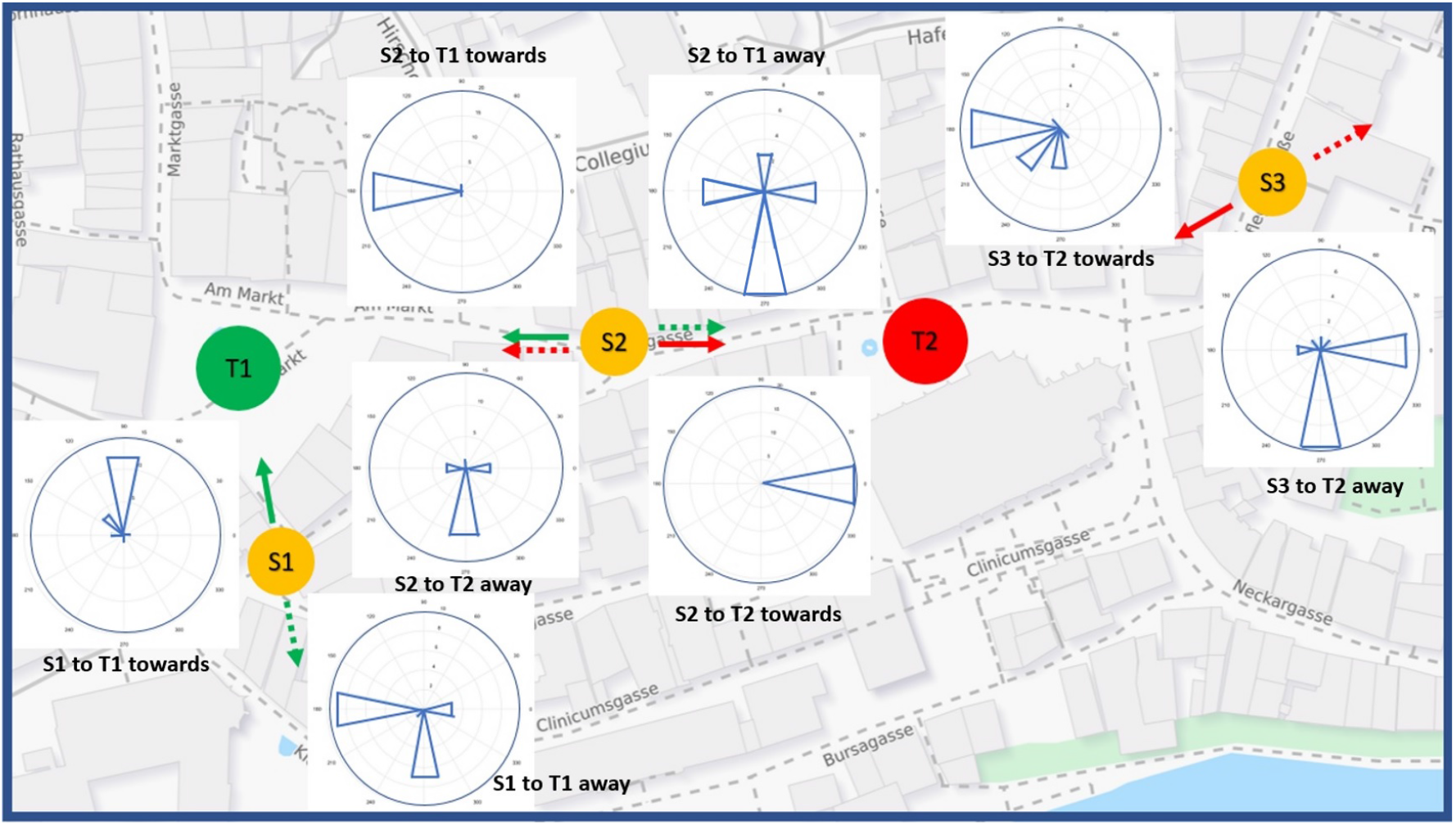
Results of Expt. 1. Sketching and target locations as in Figure 3. Green and red arrows indicate sketching of T1 and T2, solid and dashed arrows indicate towards and away conditions. Next to each arrow the polar plot of the distribution of orientations of the drawn maps is shown (north is up). Each histogram shows data from 20 sketch maps. Map source: www.openstreetmap.org

For the statistical analyses, the orientation ratings were binned into four classes north, east, south, and west. The rare intercardinal ratings were counted es 0.5 for each of the two adjacent cardinal directions. Independence of map orientation and body orientation was tested with the *χ*^2^-test. Orientation frequencies for the four task types S1-T1, S2-T1, S2-T2, and S3-T2 were concatenated into lists with 4 *×* 4 = 16 elements, separately for the towards and away conditions. The resulting *χ*^2^ statistics has 12 degrees of freedom, since the total probabilities within each task will add to unity. Data show a significant effect of body orientation (towards vs. away), *χ*^2^(12, *N* = 160) = 72.6, *p* <.001.

In addition, the imagined orientations of the target locations differ significantly between work locations. For this analysis, we concatenated the towards and away cases for task S1-T1 into an eight-bin histogram and compared it to the same histogram for the S2-T1 task. As before, the total probabilities for the four “towards” and the four “away” cases add to unity, resulting in a *χ*^2^ statistics with 6 degrees of freedom. The orientation of the sketch maps produced at S1 was found to be significantly different from the orientation of the sketch maps produced at S2: *χ*^2^(6, *N* = 80) = 25.7, *p* <.001. The same analysis was also applied to target T2 and yielded the respective result: The orientation of the sketch maps produced at S2 is significantly different from the orientation of the sketch maps produced at location S3: *χ*^2^(6, *N* = 80) = 39.8, *p* <.001.

In summary, sketch map orientation depended (at least) on two factors: sketching location and body orientation.

A further analysis of the data concerns the preferred orientations of the produced sketch maps in relation to three theoretical angles, (i) the airline direction from the sketching location to the target, (ii) the route direction heading into the street that most quickly leads from the sketching location to the target, and (iii) the canonical view of the target as indicated by the black arrows in Figure 3. Note that the route direction (ii) is also the body orientation of the subjects in the towards condition. Figure 6 shows the differences of the sketch map orientation and each of the three theoretical angles for all sketching tasks. Thus, if subjects would exactly align their maps with the airline direction, Figure 6a would show an ideal peak at angular difference 0. Significant alignment with the theoretical angles was tested with the circular *V* -test (e.g., Berens, 2009). Results show significant effects of all three theoretical angles in the towards condition (airline: *V* (80) = 64.4, *p* <.001; route: *V* (80) = 60.8, *p* <.001, canonical: *V* (80) = 27.5, *p* <.001)), while in the away condition, significant map alignment was found only with the canonical viewing direction (*V* (80) = 24.0, *p* <.001). Note that for the S2-T1 task, all theoretical angles are virtually the same.

**Figure 6:**
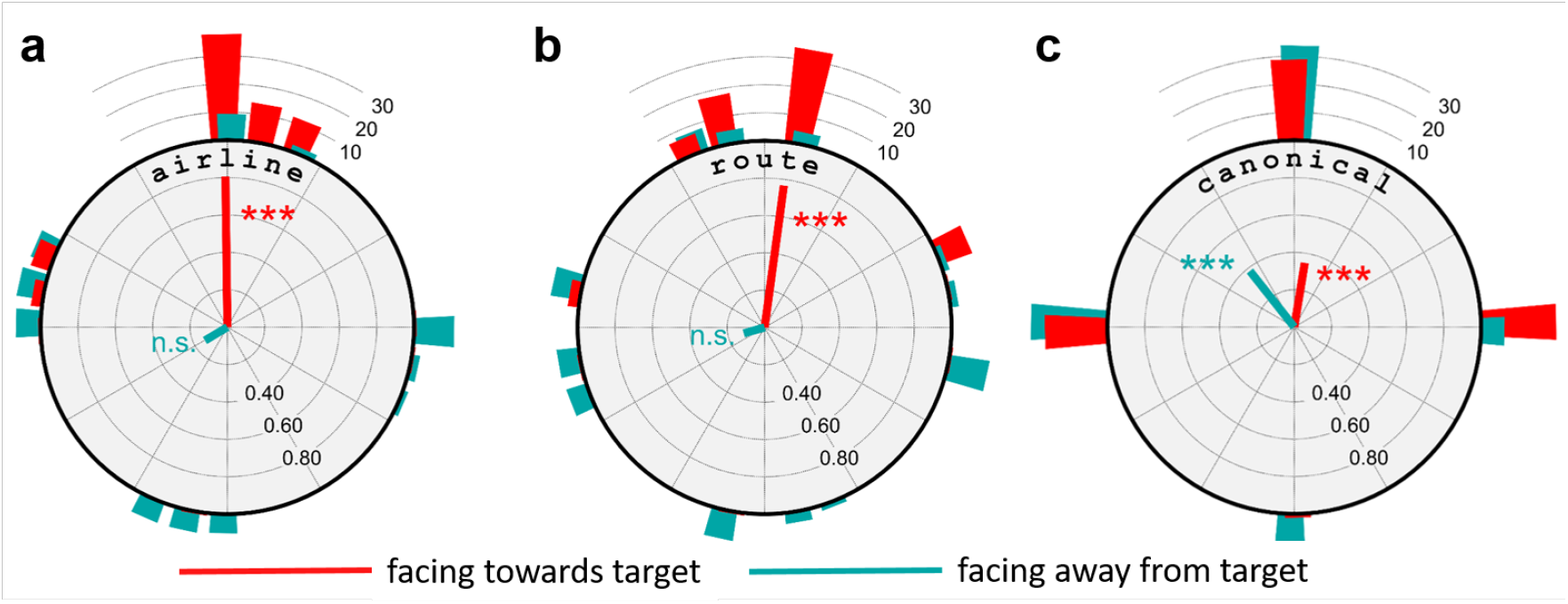
Deviation of sketch map orientation from the theoretical directions airline to goal (**a.**), route (**b.**), and canonical view (**c.**). red: towards condition, blue: away condition. Same data as in Figure 5, accumulated over all tasks. Radial lines are circular means, histogram bars show frequency data.

Results of experiment 2 appear in Figures 7 and 8. The possible effect of immersion was analyzed by concatenating the four NESW-histograms of each phase of the experiment into 16-bin lists and fusing empty bins with adjacent occupied ones. This was necessary for one bin, which reduces the number of degrees of freedom by one (from 12 to 11). The resulting test revealed a significant effect of immersion, *χ*^2^(11, *N* = 80) = 20.6, *p* <.04.

**Figure 7:**
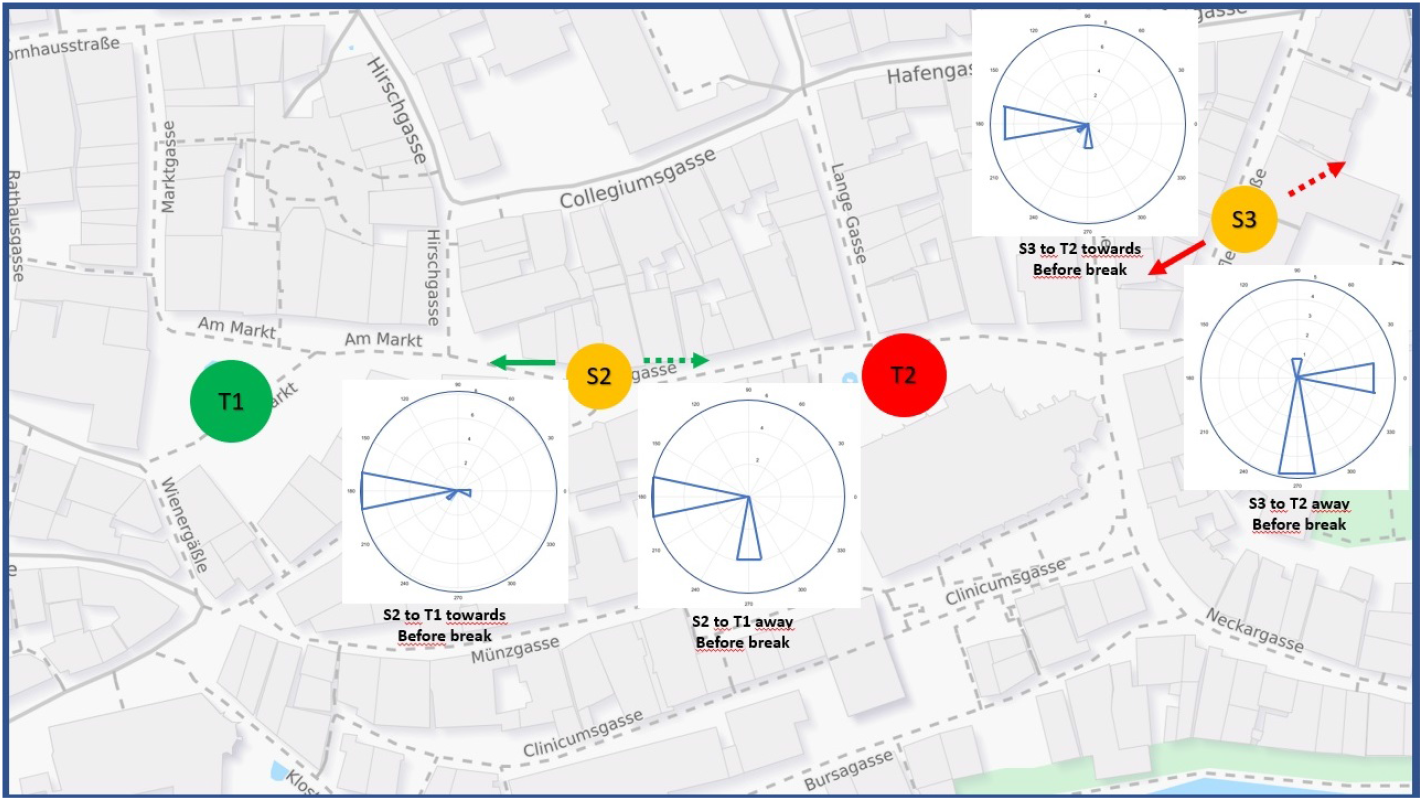
Results of the first (immersed) phase of experiment 2; for explanations see Figure 5. Map source: www.openstreetmap.org

**Figure 8:**
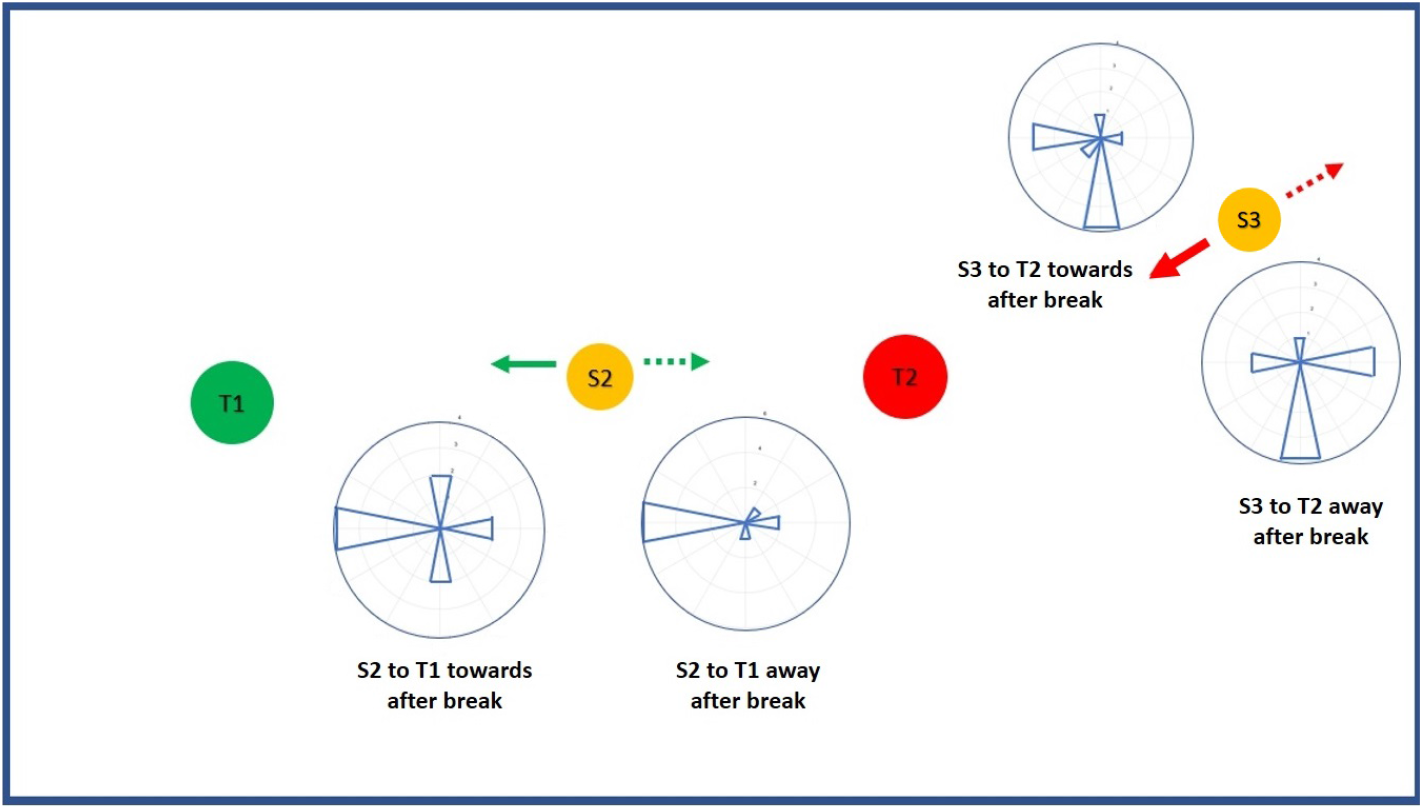
Results of the second phase of experiment 2 (after the break); for explanations see Figure 5.

Experiment 2 also reproduced the effect of body orientation demonstrated already in Experiment 1, but only for the immersed phase before the break. Again, empty bins were fused with adjacent occupied ones and the number of degrees of freedom was reduced accordingly. The orientation of the sketch maps produced before the break depended on the subjects’ body orientation (towards vs. away), *χ*^2^(5, *N* = 40) = 13.5, *p* <.05, while no such dependence was found in the second phase of the experiment after the break, when subjects produce their sketch maps in an office environment, *χ*^2^(6, *N* = 40) = 5.6, *p* >.05.

Figure 9 shows the alignment of the chosen view orientations with the theoretical directions airline to goal, route to goal, and canonical view, each with and without immersion in the virtual environment. While immersed in the virtual envirnment, highly significant alignment was found with all three theoretical directions (airline: *V* (40) = 30.8, *p* <.001; route: *V* (40) = 24.4, *p* <.001; canonical: *V* (40) = 20.0, *p* <.001). This pattern is unchanged after the break, when subjects repeat their tasks in an office environment, although the significances are weaker (airline: *V* (40) = 9.6, *p* <.05; route: *V* (40) = 11.9, *p* <.01; canonical: *V* (40) = 12.0, *p* <.01). Together with the significant effect of immersion revealed in the *χ*^2^-analysis, this indicates a weakening of orientational preference after the break without a change in the preferred orientation itself.

**Figure 9:**
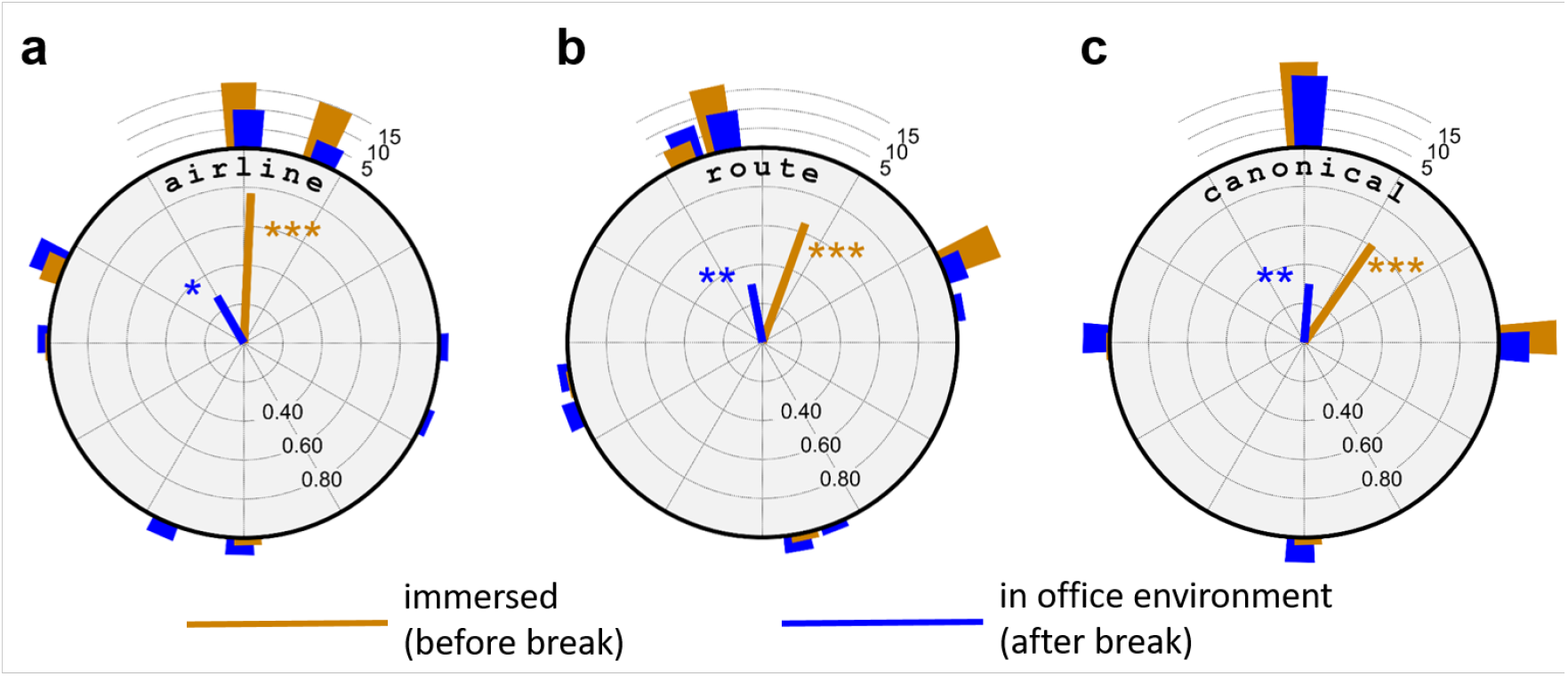
Deviation of sketch map orientation from the theoretical directions airline to goal, route to goal, and canonical view. Same data as in Figures 7 and 8, accummulated over all tasks and body orientations. Colors indicate immersion condition. Radial lines are circular means, histogram bars show frequency data.

## 5 Discussion

The results presented in this paper were obtained in a virtual environment but confirm and extend earlier findings on position dependent recall in a real urban environment by Röhrich et al. (2014). Subjects asked to produce from memory sketch maps of distant target places align their productions with specific viewing directions. These depend on the airline or route direction from the sketching site to the target, the body orientation of the subjects at the sketching location, and a standard or canonical view of the target which is also activated when the distance to the target is large (Röhrich et al., 2014). The canonical view is “allocentric” in that it does not change as the observer moves around while the airline and route axes are of course dependent on observer position, i.e. egocentric (see Klatzky, 1998). In our experiments, the airline direction was roughly identical with the direction of the shortest route connection to the target; we can therefore not distinguish the effects of these two factors.

If the subjects’ body orientation is aligned with the route direction to the target, sketch maps are oriented as if the subjects could look through the intervening buildings right to the target (“towards” cases in Figure 5). One possible interpretation of this result is that in order to solve the task subjects perform a mental travel to the target during which they imagine oriented views along the route, and sketch the final view imagined upon arrival at the target (see Basten et al., 2012). In the away condition, sketch maps are preferably aligned with the target’s canonical view (Figure 6f), i.e. the view produced also at large distances (Röhrich et al., 2014).

The difference between the sketch orientations in the towards and away conditions implies that subjects have at least implicit knowledge of their current body orientation relative to the target. This cannot be a result of path integration (as in the sensorimotor alignment effect of Kelly et al. (2007)) since subjects were teleported between sketching locations. Rather, it must be obtained from spatial memory. Memory structures suited to provide this type of information are the view-graph suggested by Schölkopf and Mallot (1995) and Röhrich et al. (2014) as well as the network of reference frames of Meilinger (2008). In both cases, the orientation changes stored for each view or reference frame transition would have to be accumulated along a path connecting the sketching and target locations. Alternatively, local orientation relative to a global reference direction could be memorized for each location (Mallot et al., 2020). Of course, information about the egocentric bearing of the target location from a sketching site would also be available from a complete metric map. In this case, however, position dependent recall itself would be hard to explain.

In the original study by Röhrich et al. (2014), body orientation was not controlled and varied substantially among subjects. The fact that the strength of position dependent recall was intermediate between the strong and absent effects found in the towards and away conditions in the present study may thus be a result of averaging over subjects with different body orientations.

One important result of this study is the overall equivalence of the real world experiments by Röhrich et al. (2014) and our immersive sketching task in a virtual environment. The role of immersion was further addressed in experiment 2 which used the same paradigm as experiment 1 but added a second phase in which subjects were retested outside the virtual environment in an office setting. Subjects do not simply reproduce their previous drawings but still show the same pattern of orientational preferences, albeit with a higher level of noise. Immersion in the virtual environment thus seems to be sufficient, but not necessary to induce position dependent recall. This in line with the findings by Basten et al. (2012) who induced position dependent recall simply by using an imagined travel as a prime.

It is interesting to compare the various angles discussed in position dependent recall with those studied in the literature on judgments of relative direction (JRD). The intrinsic axis of Shelton and McNamara (2001), i.e. the viewing direction in which an environment is most easily imagined, is a property of this environment. It does not change if the observer moves and is therefore allocentric in the sense of Klatzky (1998). The instrinisc axis is thus similar to our canonical view direction of the target place. The imagined heading direction is defined in the JRD task by the instruction to imagine facing towards a particular object. In our experiment no such instruction is used, but subjects may interpret the sketching task in a similar way. If they did, imagined heading would be the equivalent of our airline direction. It can, however, not be obtained from the imagery itself, but needs to take into account the current location and its remembered spatial relation to the target. Finally, sensorimotor alignment (Kelly et al., 2007) does not seem to play a role in our virtual environment experiments where subjects are teleported between sketching sites. Our angle of body orientation is defined by visual cues from the environment and knowledge of the spatial relations between sketching site and target. In real world experiments, this may be combined with cues from path integration.

In conclusion, position dependent recall is consistent with the idea that spatial longterm memory is organized as a graph of local places, views, or reference frames which can be “loaded” into a chart-like working memory stage for purposes of imagery and spatial planning. The representation in the chart has a specific orientation leading to view-like productions in sketching tasks. The orientations depend on the current situation and task, presumably in ways useful for spatial planning.

## Acknowledgement

The research reported in this paper was carried out at the Department of Biology of the University of Tübingen. Additional support was provided by the Deutsche Forschungsgemeinschaft (DFG) within the research unit FOR2718, Modal and Amodal Cognition, under grant no MA1038/15-1.

